# Odor Coding of Nestmate Recognition in the Eusocial Ant *Camponotus floridanus*

**DOI:** 10.1101/614719

**Authors:** S.T. Ferguson, K.Y. Park, A. Ruff, I. Bakis, L.J. Zwiebel

**Affiliations:** Department of Biological Sciences, Vanderbilt University, 465 21^st^ Avenue South, Nashville, TN 37235, USA

**Author notes:** These authors contributed equally to this work (K.Y. Park, A. Ruff).

**Keywords:** nestmate recognition, odorant receptors, orco, aggression, odor coding

## Abstract

**Background:** In eusocial ants, aggressive behaviors require a sophisticated ability to detect and discriminate between chemical signatures such as cuticular hydrocarbons that distinguish nestmate friends from non-nestmate foes. It has been suggested that a mismatch between a chemical signature (label) and the internal, neuronal representation of the colony odor (template) leads to the recognition of and subsequent aggression between non-nestmates. While several studies have demonstrated that ant chemosensory systems, most notably olfaction, are largely responsible for the decoding of these chemical signatures, a definitive demonstration that odorant receptors are responsible for the detection and processing of the pheromonal signals that regulate nestmate recognition has thus far been lacking. To address this, we have developed an aggression-based bioassay incorporating a suite of highly selective odorant receptor modulators to characterize the role of olfaction in nestmate recognition in the formicine ant *Camponotus floridanus*.

**Results:** Validation of our aggression-based behavioral assay was carried out by demonstrating an antennal requirement for nestmate recognition. In order to adapt this bioassay for the volatile delivery of Orco modulators, electroantennography was used to show that both a volatilized Orco antagonist (VUANT1) and an Orco agonist (VUAA4) eliminated or otherwise interfered with the electrophysiological responses to the hydrocarbon decane, respectively. Volatilize administration of these compounds to adult workers significantly reduced aggression between non-nestmates without altering aggression levels between nestmates but did not alter aggressive responses towards a mechanical stimulus.

**Conclusions:** Our studies provide direct evidence that the antennae (as olfactory appendages) and odorant receptors (at the molecular level) are necessary for mediating aggression towards non-nestmates. Furthermore, our observations support a hypothesis in which rejection of non-nestmates depends on the precise detection and decoding of chemical signatures present on non-nestmates as opposed to the absence of any information or the active acceptance of familiar signatures. In addition to describing a novel approach to assess olfactory signaling in genetically intractable insect systems, these studies contribute to a long-standing interest in odor coding and the molecular neuroethology of nestmate recognition.

## Background

Aggression comprises a range of biologically salient social interactions with implications for individual behavior as well as the collective integrity of animal societies. While hostile behaviors can be observed throughout the Metazoa (1-5), recently established experimentally tractable eusocial insect models present an opportunity to investigate the mechanistic basis of aggression within a social context. In this regard, ants provide a compelling model for the study of aggression and its triggering mechanisms. Ant colonial lifestyles and reproductive hierarchies are maintained by aggressive social interactions that are modulated by their ability to detect, discriminate between, and respond to a large array of chemical cues often known as pheromones (2, 6-8). Moreover, recent studies (9, 10) have demonstrated the value of applying novel genetic and molecular techniques that have restricted availability/utility in the study of humans and other social primates.

The formicine ant *Camponotus floridanus* live in colonies that are founded by a single reproductive queen (2, 11). Workers nurse the queen’s offspring, forage for food, and defend nest and territory from non-nestmates (nNMs) (2). Although individual workers contribute to broader colony-level phenotypes, the integrity of social behaviors depends on the collective actions of the colony (12). Among these social behaviors, nestmate (NM) recognition is especially important for establishing and maintaining discrete societal boundaries for *C. floridanus* and many other species of ant (2). NM recognition is a dynamic behavior that has been postulated to occur when an individual ant compares chemically encoded “labels” that it encounters with potentially multiple neural-encoded “templates” that represent its own particular global colony chemosensory signature whereby a mismatch between a foreign label and the recognition templates leads to aggression between nNMs (13-15). The foreign label is derived, at least in part, from subtle variations in the profile of cuticular hydrocarbons (CHCs) that distinguish nNMs from NMs (6, 13, 16).

Early genetic models provided a framework for understanding the criteria required to assess colony membership status when comparing the recognition template to a respective label (17). These have been broadly organized into two categories: the gestalt model, in which label sharing between individuals yields a distinct template based on a blend; and individualistic models, which include requiring the exact matching of the label to the template (“genotype matching”), rejection of any labels containing cues not found in the template (“foreign-label rejection”), and the acceptance of labels that overlap with the template (“habituated-label acceptance”). Similarly, there have been efforts to elucidate the rules governing template-label matching within a phenotypic context (13, 16, 18). These models suggest that ants discriminate between friends and foes based on the presence and/or absence of NM (“desirable”) cues or nNM (“undesirable”) cues. While it was initially proposed that ants accept individuals if they possess desirable cues (D-present) or if they lack undesirable cues (U-absent) to the exclusion of all others (18), more recent evidence suggests that ants actively detect foes but not friends through the detection of nNM odor cues (simple U-present model) (16). Importantly however, discrimination may also occur when critical components of the CHC profile are missing (13). These studies suggest that there are multiple templates being used to assess different labels, and that there is variability in the importance of a given component of the label, whether in absence or in abundance, when determining nNM or NM status.

While the importance of CHCs in mediating NM recognition among ants is well established, several alternative hypotheses have been proposed for the neuronal and molecular mechanisms required for ants to distinguish friends from foes (13, 16-21). In all of these models, CHCs and other semiochemicals are initially detected by the peripheral olfactory sensory system which relies on three major classes of peripheral chemosensory receptors–odorant receptors (ORs), gustatory receptors, and ionotropic receptors. Insect ORs are expressed in olfactory receptor neurons (ORNs) housed within sensilla on the antennae (reviewed in (22)), where they function as heteromeric complexes consisting of an obligate and conserved OR co-receptor (Orco) and at least one “tuning” OR that determines odorant (ligand) specificity (23-29). Several studies have revealed a large expansion of the *OR* gene family in ants as well as other eusocial insects (23, 30-37), leading to the demonstration that this expanded chemoreceptor family is responsible for the detection of socially relevant chemical cues such as CHCs (38, 39).

Despite the long-held appreciation for the role of CHCs and other chemical cues in mediating NM recognition and social behaviors in ants, little is known about the specific molecular components of olfactory signal transduction that are active in regulating NM recognition and the triggering of aggression toward nNMs. Electrophysiological studies of *Camponotus japonicus* first suggested that a dedicated multiporous NM recognition sensilla exhibited an all-or-none response to nNM CHC blends but, importantly, did not respond to NM CHC blends–thus leading to a model in which ants are desensitized and ultimately anosmic to their own odor cues (19). In contrast, recent studies using both antennal electrophysiology and antennal lobe calcium imaging in the related ant species *C. floridanus* demonstrate these ants are capable of detecting both nNM and NM odors (20, 21, 40). It has been proposed that these seemingly contradictory findings support a model in which two sensilla subtypes—one broadly tuned to hydrocarbons and the other tuned to specific hydrocarbons—facilitate habituation to different labels (41).

The paucity of data in this regard may be attributed, at least in part, to the challenges of targeted molecular approaches currently available in the study of Hymenopteran insects. The development of these techniques represents an important step towards understanding the function and evolution of the molecular mechanisms involved in complex social behaviors such as NM recognition with the potential to shed light on longstanding questions within the field of social insect biology. To begin to address this, a series of behavioral, physiological, and gene knockout studies were carried out to characterize the relationship between ant ORs and CHCs as well as other biologically salient chemical cues. These studies demonstrated that CHCs and other general odorants were broadly detected across the various OR subclades while CRISPR-mediated gene knockout of *orco* resulted in alterations of both solitary and social behaviors as well as profound neuroanatomical disruptions in the antennal lobe (9, 10, 38, 39). Taken together, these studies suggest that ORs play a critical role not only in a diversity of behaviors but also importantly in ant neural development.

Here, we report studies that specifically address the odor coding of NM recognition by utilizing a novel volatilization paradigm incorporating a set of highly selective Orco agonists and antagonists to acutely and globally impact OR-based pathways in the context of an aggression bioassay. In this manner, we are able to directly test the hypotheses that aggression is triggered by the active detection and decoding of discrete chemosensory stimuli and more specifically that the functionality of the OR-Orco ion channel complex is necessary for NM recognition.

## Results

### Nestmate Recognition Requires Antennal-based Signaling

We first aimed to develop an olfactory-based NM recognition bioassay in which two ants—NMs from the same home colony or nNMs from two different colonies—were able to interact with one another after an acclimation period (Fig. 1A). To this end, we initially took a broad approach to assess the role of olfactory signaling in modulating NM/nNM aggression in the context of pairwise trials conducted using adult *C. floridanus* minor worker ants with either unilateral or bilateral antennal ablations. In these studies, both control *C. floridanus* workers as well as those having undergone unilateral ablations were able to routinely discriminate nNMs from NMs and display only nNM aggression (Fig. 1B). In contrast, ants with bilateral antennal ablations displayed a significant and indeed near-complete reduction in aggression against nNMs. These data are consistent with the widely reported ability of *C. floridanus* workers to robustly discriminate between nNMs and NMs and supports the hypothesis that their chemosensory apparatus is required to recognize and trigger aggression against nNMs (2, 6, 13, 16, 19, 20, 38, 39, 42-44).

**Fig. 1.**
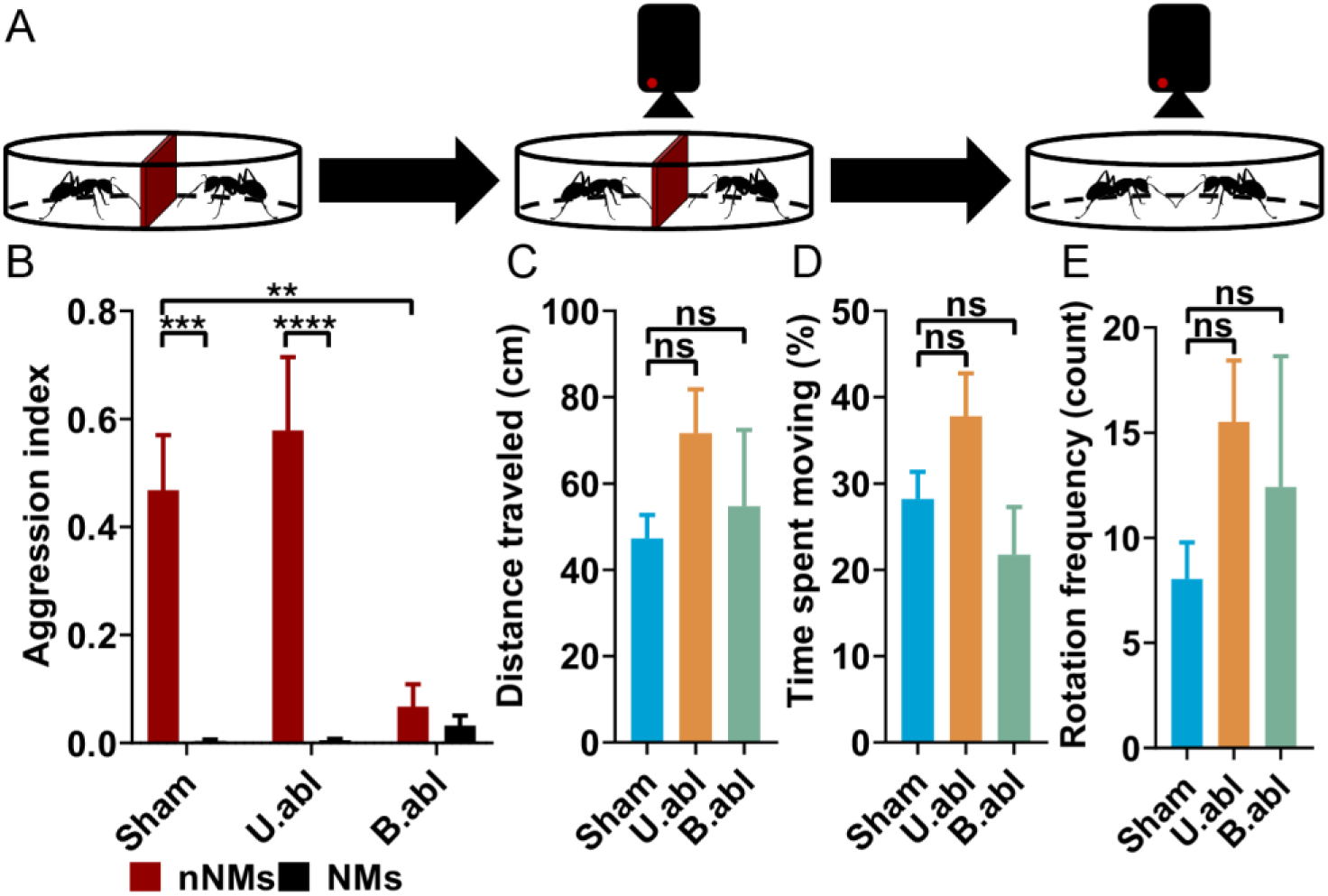
Aggression and mobility responses of adult minor workers following antennal ablation (Sham = control; U.abl = unilateral ablation; B.abl = bilateral ablation). (**A**) Schematic of the ablation bioassay depicting the acclimation period (left), mobility controls (center), and aggression bioassay (right). (**B)** Bilateral antennal ablations significantly reduce nNM aggression compared to the sham control (Two-Way ANOVA, N=6-10). (**C-E**) There is no significant difference across the mobility parameters tested between the sham control and either of the ablation treatments (Kruskal-Wallis Test, N=24-29). Error bars display S.E.M. Asterisks indicate P-value: **<0.01, ***<0.001, ****<0.0001.

To further control for potentially confounding variables—including the outright death or incapacitation of the ants due to the damage sustained from the ablations—we measured a number of other behavioral indicators including total distance traveled, percentage of time spent moving/not moving, and the frequency of rotations using an automated tracking program (see Methods). Here, the activity of a single ant was recorded for three minutes immediately following the 10-minute acclimation period and preceding the ablation aggression bioassays (Fig. 1A). These assays revealed no significant difference between the sham control and either of the ablation treatments (Fig. 1C-E). That treated ants were able to recover from the injury and retain fundamental aspects of mobility coupled with the observation that unilaterally ablated workers maintained the ability to discriminate between NMs and nNMs suggests that the decrease in aggression was likely due to the absence of antennae-mediated signaling as opposed to confounding variables introduced by the ablation treatment. However, as the removal of the antennae disrupts a broad range of both mechanoreceptors as well as chemoreceptors (45), a more targeted approach is required to assess the specific function of OR-dependent chemoreceptor signaling in this context.

### Nestmate Recognition is an Active, OR-dependent Process

In order to further examine this process within the narrow context of assessing the role of ORs in NM recognition and aggression, we adapted our bioassay to incorporate the sustained volatile administration of a set of highly specific Orco allosteric modulators (Fig. 2A, Additional File 1). The first member of this unique class of pharmacological agents (known as VUAA-class actives) was initially identified through high-throughput screening for small molecule activators of Orco/OR complexes expressed in HEK293 cells (28, 29, 46). In subsequent studies that revealed extraordinarily narrow structure-activity relationships, several additional VUAA-class actives were identified and characterized that now comprise several more potent agonists (including VUAA4 used here), a non-competitive antagonist (VUANT1, used here) as well as an inactive structural analog (VUAA0, used here) (28, 46-49). Further studies, including single-sensillum recordings of female-specific basiconic sensilla in *C. floridanus*, have demonstrated that the potency of these modulators in both volatile and non-volatile form is conserved across a wide range of insect orders (40, 47, 50-52). Indeed, VUAA-Orco interactions have recently been directly confirmed by cryo-electron microscopy studies characterizing the structure of an Orco tetramer from the parasitic fig wasp *Apocrypta bakeri* (53).

**Fig. 2.**
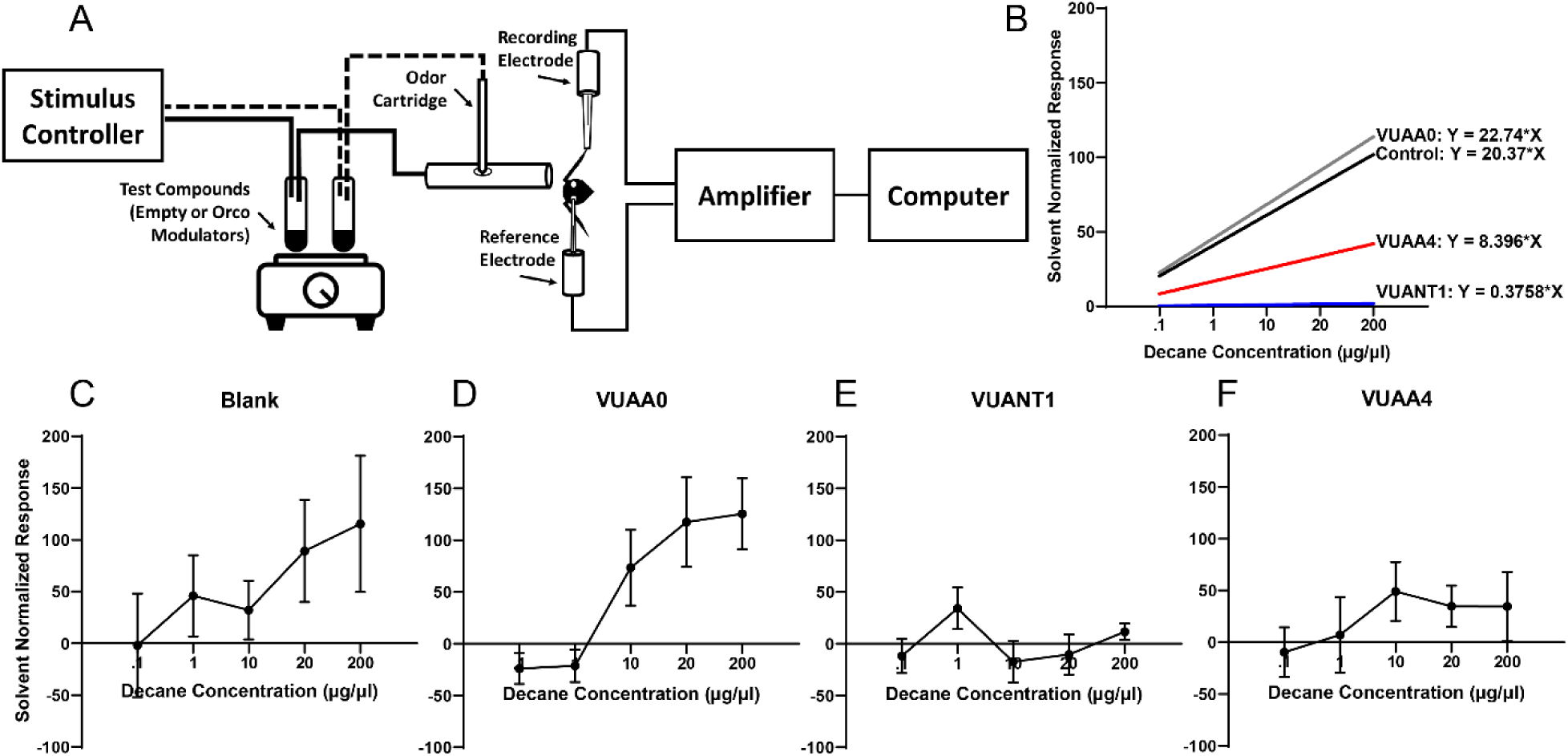
Electrophysiological responses of adult minor workers to the hydrocarbon decane under different background airflow conditions (Blank = heated air alone; VUAA0 = inert chemical analog control; VUANT1 = Orco antagonist; VUAA4 = Orco agonist). (**A**) Schematic of the electroantennograms. (**B**) Best-fit lines derived from the solvent (hexane) normalized responses to serial concentrations of decane for Blank (**C**), VUAA0 (**D**), VUANT1 (**E**), and VUAA4 (**F**) backgrounds. The slope of the best-fit line for Blank, VUAA0, and VUAA4 are significantly different from 0 (Linear Regression, N=5-6, see Additional File 3). Error bars display S.E.M.

The use of these unique and highly specific chemical tools allows us to selectively target Orco and therefore the functionality of all OR/Orco complexes to examine NM recognition with altered OR signaling in otherwise wild-type adult *C. floridanus* workers. This is an essential aspect of our approach in light of the broad neuroanatomical alterations that have recently been observed in the development of the antennal lobes of Orco mutants in two ant species (9, 10) which are reasonably likely to impact olfactory processing. Indeed, the use of volatile Orco modulators represent a novel and requisite approach for disrupting OR functionality in insects such as ants that require alternatives to CRISPR-mediated targeting of pleiotropic genes such as *orco* (9, 10).

In order to validate the efficacy of the VUAA-class actives delivered within a constant background airflow to our aggression bioassay arena, we performed electroantennograms (EAGs) to assess whole antennal responses to several concentrations of the hydrocarbon decane (C10) in adult workers exposed to heated air (blank control) or volatilized compound (Fig. 2A). Here, we observed similar dose-dependent responses in both our blank control and VUAA0 (Fig. 2B-D). Indeed, linear regression analysis revealed that the slope of the blank control and VUAA0 are significantly different from 0 (i.e. a flat line) (Fig. 2B, Additional File 3). Consistent with expectations, the slope of VUANT1 is not significantly different than 0 (Fig. 2B and E, Additional File 3), suggesting that exposure to this compound completely eliminated dose-dependent detection of decane. While volatile administration of VUAA4 also clearly disrupts hydrocarbon detection, it results in an intermediate phenotype, displaying a muted and partially dose-dependent response with seemingly static, yet low, responsiveness at higher concentrations (Fig. 2B and F). These are likely the result of broad ORN desensitization after prolonged exposure to this potent Orco agonist. Nevertheless, the slope of VUAA4 is significantly different from 0 (Fig. 2B, Additional File 3), suggesting that dose-dependent hydrocarbon detection and ORN firing still occur albeit not in the same manner as the controls. Taken together, these results suggest that acute volatile administration of VUAA-class actives can indeed be used to disrupt Orco-mediated olfactory signal transduction in ants.

Using this newly established volatilization paradigm, we next sought to determine the precise role of OR-signaling in mediating aggression towards nNMs. Ants taken from across nine independent colonies exposed to either Orco modulator displayed a significant reduction, and indeed a near complete elimination, of aggression towards nNMs (Fig. 3A). Importantly, in addition to the inability to aggressively respond to nNMs, ants treated with either the Orco agonist or the antagonist displayed no alteration in their non-aggressive responses to NMs. This lack of misdirected aggression toward NMs as well as the failure to correctly attack nNMs in ants treated with these highly selective Orco/OR modulators demonstrates that, in *C. floridanus*, aggression is specifically mediated by the OR-dependent detection of specific and unambiguous odor cue signatures from nNM foes rather than the general absence or incorrect processing of familiar signatures of NM friends.

**Fig. 3.**
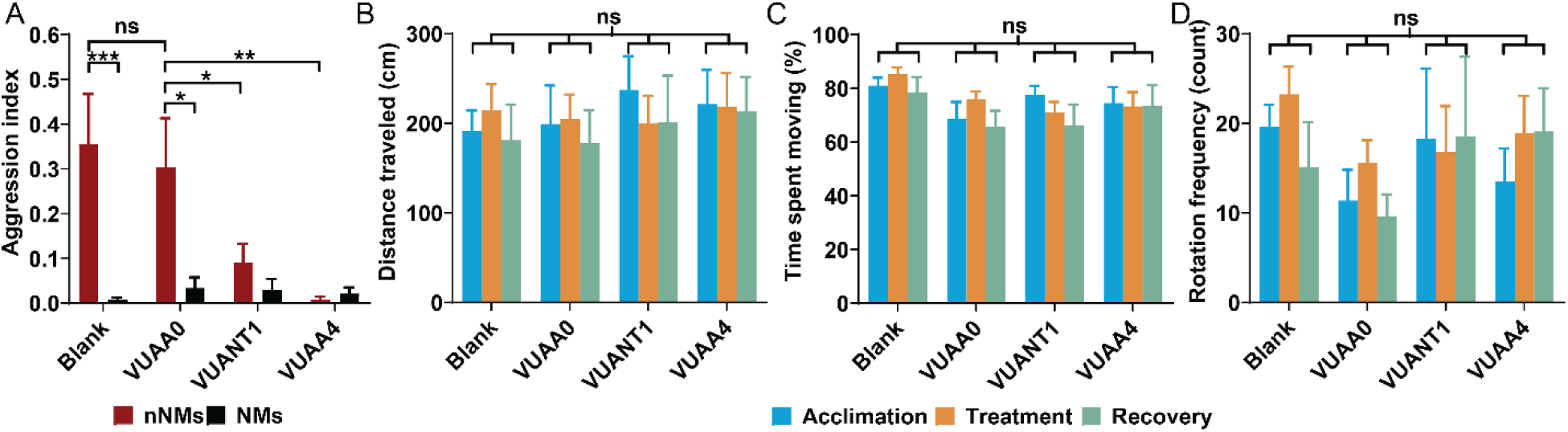
Aggression and mobility responses of adult minor workers during exposure to volatilization treatments (Blank = heated air alone; VUAA0 = inert chemical analog control; VUANT1 = Orco antagonist; VUAA4 = Orco agonist). (**A**) Disrupting Orco-mediated olfactory signal transduction significantly reduces aggression towards nNMs (Two-Way ANOVA, N=10-12). (**B-D**) There is no significant interaction between treatments across the mobility parameters tested (RM Two-Way ANOVA, N=7-9). Error bars display S.E.M. Asterisks indicate P-value: *<0.05, **<0.01, ***<0.001.

Furthermore, in order to assess whether the disruption of OR-signaling reduces aggression within the narrow social context of NM recognition or alternatively acts to broadly inhibit aggressive behaviors, we conducted parallel bioassays that utilized mechanical rather than chemical stimuli to evoke aggression. Here, using a modified aggression bioassay based on previous methods described in (54) and (55), individual ants were challenged with a chemically neutral mechanical stimulus (i.e. a clean Von Frey filament) and subsequently scored for biting responses as well as wide opening of the mandibles as indicators of aggression. Importantly, inasmuch as there was no significant difference in aggression among the various treatment groups (Additional File 2), we can conclude that disrupting Orco-mediated olfactory signaling does not generally inhibit aggressive responses in *C. floridanus* but instead specifically impacts workers’ ability to discriminate NMs from nNMs and aggressively respond to the latter.

In order to further control for potentially confounding variables in response to these volatilization treatments, the activity of a single ant was recorded immediately following a 10-minute acclimation period. These trials consisted of a continuous 9-minute bioassay separated into three 3-minute segments. During the first segment, the ants were exposed to a continuous flow of untreated air (‘Acclimation’); for the second segment, the ants were exposed to a continuous flow of volatilized VUAA-class active or untreated air in the case of the blank control using the same parameters established for the volatilization aggression bioassay (‘Treatment’); and lastly, during the third segment, the ants were again exposed to a continuous flow of untreated air (‘Recovery’). A Y-junction connected to the compressed air tank alternated between the empty test tube during the Acclimation and Recovery phases and the treatment or blank tube during the Treatment phase. An examination of overall mobility parameters revealed no significant interaction effect when comparing control ants and ants treated with either an Orco agonist or antagonist before, during, or after exposure to each treatment (Fig. 3B-D).

## Discussion

In ants and other eusocial insects, NM recognition depends on the ability to discriminate between self and non-self where the recognition of non-self—in this instance nNMs—often leads to aggression (reviewed in (56)). While it is clear that these aggressive responses are mediated by the detection of subtle differences in the CHC profiles that demarcate individual colonies (6, 13, 16, 42), the precise coding of that information within the olfactory system has remained ambiguous and, to some extent, controversial. Initially, we took a conservative approach to validate both our bioassay along with the expected antennal requirement (43, 44) for NM recognition (Fig 1). Once established, this experimental paradigm was further adapted to accommodate the sustained volatile administration of highly specific VUAA-class Orco modulators to directly test the hypothesis that NM recognition in adult *C. floridanus* workers is solely dependent upon OR-based olfactory signaling as well as facilitate the characterization of odor coding in this process. In light of the broad developmental defects that result from the loss of Orco in other ant systems (9, 10), these pharmacological tools provide a unique opportunity to acutely examine the role of OR-based signaling in an otherwise wild-type adult nervous system. At the same time, in light of the obligate colocalization of Orco together with tuning ORs in every insect ORN (24, 48, 57), exposure to Orco modulators is expected to have profound and widespread effects.

As previously observed in other contexts (40), treatment with the VUANT1 antagonist effectively silences all Orco/OR complexes and prevents the generation of any interpretable signal (Figure 2). In the case of the VUAA4 Orco agonist, activation of all Orco/OR complexes leads to broad ORN desensitization resulting in significantly diminished signaling (Figure 2) that we postulate effectively generates an uninterpretable or “confused” coding signal. In either case, the lack of any odor signal or the presence of imprecise odor cues that are expected after treatment with an Orco antagonist or agonist, respectively, are both equally insufficient to elicit aggression between nNMs (Fig. 3).

The observation that an Orco antagonist decreases aggression between nNMs is broadly consistent with a simple U-present rejection model and supports the view that ants are not actively recognizing friends (16, 58). However, the curious finding that an Orco agonist, which would be expected to generate a foreign label different from that of the endogenous template, would also decrease aggression between nNMs rather than increase aggression between NMs suggests that the simple presence of foreign or otherwise imprecise cues are also insufficient to elicit aggression. These studies therefore support a model in which an unambiguous triggering stimulus must be precisely detected in order to evoke aggression. As such, we propose that the recognition mechanism in *C. floridanus* occurs via a lock-and-key mechanism whereby the specific parameters of the foreign chemical label key, defined by the combinatorial presence and/or absence of salient odor cues, must be precisely decoded by an OR-mediated lock (Fig. 4). Under this assumption, ants may identify nNMs in two different ways which are not necessarily mutually exclusive: 1. unfamiliar nNM labels are compared to a familiar NM template with bounded thresholds wherein the label must be sufficiently different from the template but not so different as to be ambiguous; or 2. unfamiliar nNM labels are compared to intruder templates that represent odor profiles which should be rejected from the colony and a certain level of precision between the label and template is required to elicit aggression.

**Fig. 4.**
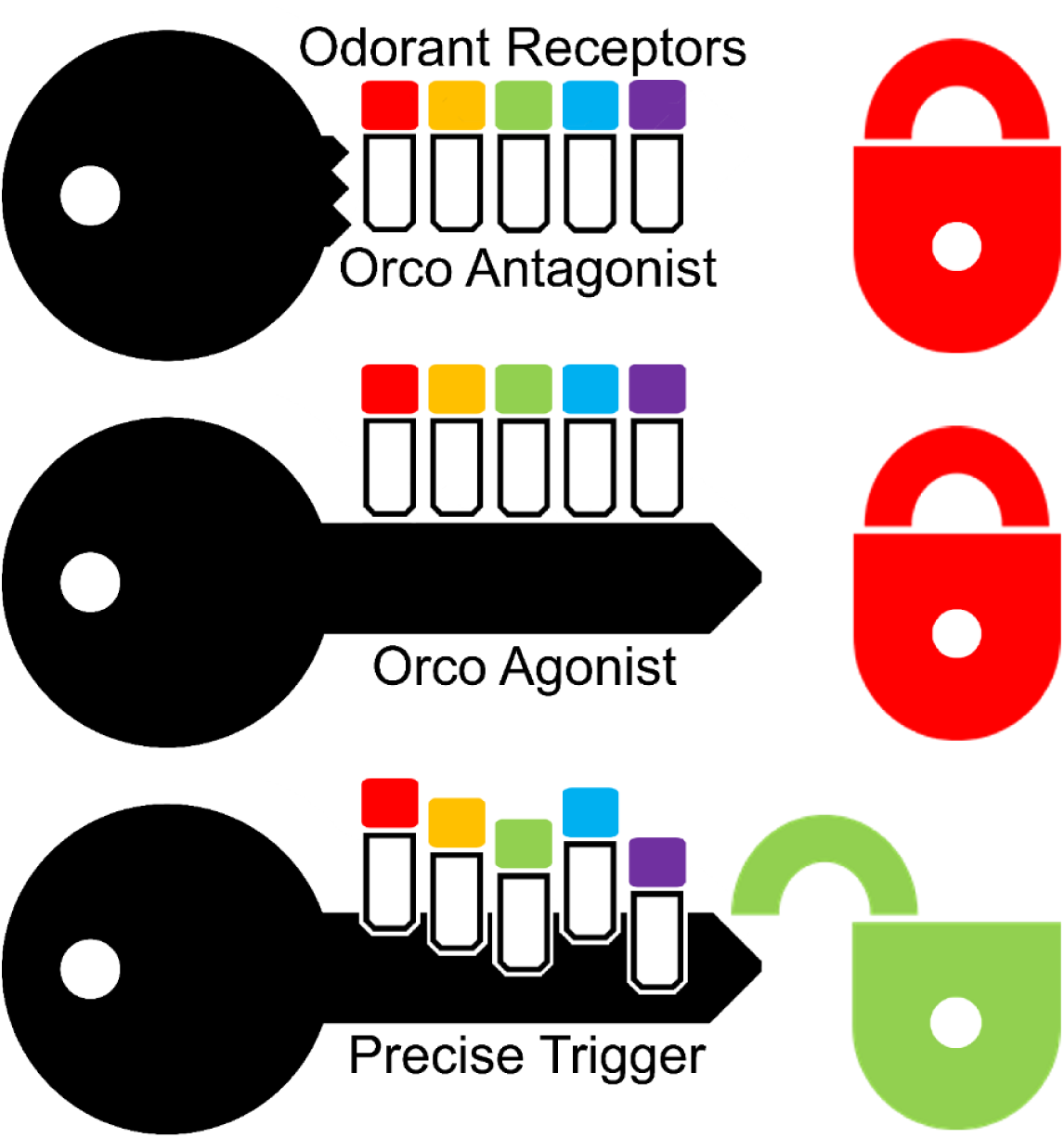
Lock-and-key model of nNM recognition and aggression. The triggering stimuli, represented by the teeth on a key, must be precisely detected by the OR-tumblers in the lock. OR-dependent recognition of nNM cues leads to aggression against foes (green open lock); however, blocking OR-dependent recognition of NM/nNM cues does not lead to aggression nor does the presence of an ambiguous chemical cue (closed red locks).

Furthermore, these data suggest that, when faced with some level of uncertainty, *C. floridanus* workers default towards acceptance rather than rejection. Over and above the benefits of conserving energy by avoiding potentially unnecessary aggression, for ants that spend the majority of their life cycles within colonies where they are more likely to encounter NMs than nNMs, this strategy may also reduce acceptance errors and therefore increase overall colony fitness (59). It will be interesting to determine whether similar processes occur across worker behavioral task groups that may spend more time outside the nest (i.e. scouts and foragers) or whether different recognition methods have evolved across castes and/or species.

## Conclusions

At a mechanistic level our data effectively excludes the sufficiency of other signaling pathways and sensory modalities and demonstrates that Orco/OR-mediated signaling is necessary for the active detection and precise processing of a discrete stimulus that triggers aggression towards nNMs in *C. floridanus*. These results are consistent with previous literature suggesting that aggression-mediated NM recognition may be more appropriately described as nNM recognition (16, 58). While the roles of individual ant ORs or even specific subsets of ORs in nNM recognition remain to be elucidated, the combinatorial interactions that are expected even among specialized ORs (38, 39), the plasticity of the potentially numerous neuronal templates (13, 42) and the similarly diverse and plastic labels (60-63) as well as the observation that even repeated stimulation with colony odors produced variable response patterns in the antennal lobe (20) are likely to make those studies extremely challenging. Nevertheless, the demonstration that precise and unambiguous OR-based coding is necessary for ants to distinguish foe from friend represents a significant advance to link the longstanding interest in social insect behavior with more recent studies detailing the evolutionary complexity of the insect olfactory system (2, 23, 30).

## Methods

### Ant Husbandry

Nine distinct laboratory colonies of *Camponotus floridanus* originating from field collections generously obtained by Dr. J. Liebig (Arizona State University) from the Long Key (D242) and Sugarloaf Key (D601) and Dr. S. Berger (University of Pennsylvania) from the Fiesta Key (C6, K17, K19, K28, K31, K34, and K39) in South Florida, USA. All colonies were independently maintained at 25°C, ambient humidity, with a 12-h light:12-h dark photoperiod. Each colony was provided with Bhatkar diet, crickets, 10% sucrose solution, and distilled water three times per week. Adult minor workers were used for all experiments and were sampled from throughout the colony.

### Ablation Aggression Bioassay

Tests were conducted during the ZT diel light cycle between ZT2 and ZT12 at ambient room temperature and humidity and performed using a six-well culture plate with polytetrafluoroethylene-coated well walls (DuPont®). Individual wells of the six-well culture plate served as distinct bioassay arenas for behavioral trials (Additional File 1). In preparation for experiments, each well (9.6cm^2^) of the six-well culture plate was fitted with a removable plastic divider that partitioned the well into two halves. The six-well culture plate and dividers were sterilized using ethanol, air dried, and positioned on top of a light box. Each individual bioassay well utilized two adult minor ants that were selected from either the same home colony (NMs) or two distinct colonies (nNMs). All ants were handled wearing gloves and using sterile, soft-tipped metal forceps and were subsequently discarded after each bioassay to ensure each ant was used only once.

Subject ants were briefly anesthetized with CO_2_ before removing their antennal flagella via an incision across the distal portion of the scape using a clean, unused razor blade. Bilaterally ablated ants had both flagella removed while unilaterally ablated ants had only a single (right or left, randomly selected) flagellum removed. Sham treated ants were anesthetized with CO_2_, and the razor was gently touched to the antennae without damaging any structures. Subsequent to ablation (or sham) treatment, ants were allowed to recover along with similarly treated NMs for at least 2 hours prior to testing.

Prior to bioassays, two ants (NMs or nNMs) were placed into each well arena, one in either half, and allowed 10 min to acclimate to handling. To document normal ant behavior within each well arena, mobility was recorded using a digital high definition camera (Panasonic® HC-V750) for 3 min (detailed below). The plastic divider within each well arena was subsequently removed and all ant interactions again recorded for 3 min. The order in which the treatments were conducted as well as the colony the ants were selected from for any given trial were randomized using RANDOM.ORG (Randomness and Integrity Services Ltd.).

### Electroantennography

Electroantennograms were performed using an IDAC-232 (Ockenfels Syntech GmbH, Germany) controller linked to a Windows XP computer running EAG2000 (Ockenfels Syntech GmbH, Germany) software. A set of 12×75mm test tubes placed atop a heat block set at 260°C containing 0.025g of the respective treatment compound (VUAA0, VUANT1, or VUAA4) or an empty tube (blank control) were connected to a Syntech CS-05 Stimulus flow Controller (Ockenfels Syntech GmbH, Germany). Using this setup, both the constant background airflow as well as the 500-ms pulse of stimulus compound contained volatilized VU-class compounds or heated air (in the case of the blank control).

Subjects ants were placed in a 20µL disposable pipet tip that was modified such that the tip opening was sufficiently wide to allow the unimpeded exposure of the head and antennae. To prevent movement of the preparation which might otherwise reduce the signal-to-noise of the recordings, the head and mandibles of the ant were restricted with wax. Borosilicate glass capillaries (FIL O.D.:1.0mm, World Precision Instruments, Inc.) were customized for EAGs on a P-2000 laser micro-pipette puller (Sutter Instruments), backfilled with 10^−1^ M KCl and 0.05% PVP buffer and placed over tungsten electrodes. A 30-guage needle was used to puncture the right eye to allow for insertion of the reference electrode. The recording electrode was placed over the distal tip of the left antenna. Decane (C10) (CAS: 124-18-5, Sigma-Aldrich) was serially diluted in hexane (0.1 µg/µl, 1 µg/µl, 10 µg/µl, 20 µg/µl, and 200 µg/µl). An odor cartridge was filled with 10µl of decane solution (or hexane alone as a solvent control) and a handheld butane torch (BernzOmatic, Worthington Industries) was used to volatilize the compound by heating the odor cartridge for 1.5 seconds. Serial concentrations were assayed sequentially starting with the lowest concentration and ending with the highest concentration. Responses were normalized to the hexane solvent control (set at 0) to account for changes in sensitivity and/or antennae degradation over time throughout the assay, and these values were used for subsequent data analysis.

### Volatile Orco Modulator Aggression Bioassay

To facilitate the administration of a continuous flow of air containing volatilized VUAA-class compounds (all custom synthesized as dry solids in-house at Vanderbilt University (28, 47-49)) into the aggression arena, bioassays were conducted in arenas consisting of modified square plastic boxes with a total area of 85cm^2^ (Pioneer Plastics Inc. ®) (Additional File 1). Mirroring the electroantennography, conditioned air (78% Nitrogen, 21% Oxygen) was delivered (at a constant 34kpa) from a compressed source (Nashville Gas LLC) to the test arena through a 12×75mm test tube atop a heat block set at 260°C which contained 0.025g of the respective treatment compound (VUAA0, VUANT1, or VUAA4) or an empty tube (Blank control) via 18G needles inserted into a rubber septum affixed to the top of the test tube before exiting through a dedicated exhaust system. Trials were recorded using a digital high definition camera and scored as described below. Although two plastic tubes were affixed to the arena during the volatilization aggression bioassays, only a single tube was actively delivering the test compound or heated air control (Additional File 2). In each assay, ants were acclimatized underneath 35mm Petri dish lids (prewashed with ethanol) for 10 minutes after which the lids were then removed (allowing the ants to interact), the airflow started, and the ants were then recorded for the 3-minute test period. All treatment compounds were randomized and coded independently such that the investigator was blinded to the treatment identity. Furthermore, the sequential order in which the compounds were tested as well as the colony the ants were selected from for any given trial was randomized using RANDOM.ORG (Randomness and Integrity Services Ltd.).

### Aggression Bioassay Scoring

Digital video recordings of all bioassays were viewed post hoc and aggression incidents manually scored for analyses. Trials in which ants did not interact, were disrupted physically during removal of the plastic barrier, or were fatally encumbered at trial onset were discarded from further analyses along with their respective mobility controls in the case of the antennal ablation bioassays. These interactions were scored by three independent, blinded observers in 10 s intervals using a binary scale such that aggression either did or did not occur (a score of 1 or 0, respectively; Additional Files 10-11). Prior to scoring, each observer was trained to recognize “aggression” as instances in which one or both ants were lunging, biting, or dragging one another. Each 10 s time interval was scored as either containing an instance of aggression or not to establish the proportion of time the ants were engaged in aggressive behavior. An aggression index was calculated by dividing the number of observed acts of aggression by the total number of observed time intervals. The mean aggression index of each video recording across all three independent scores was used for subsequent statistical analysis.

### Mobility Control Parameters

Mobility control videos were analyzed using an automated tracking software package (Ethovision® XT v8.5, Noldus Information Technology) to calculate total distance traveled (cm), percentage of time spent moving (%), and the frequency of rotations (count). Time spent moving/not moving was calculated with thresholds of 0.30cm/s (start velocity) and 0.20cm/s (stop velocity) as determined by the EthoVision® XT software with an averaging interval of 1 sample. To determine the percent of time spent moving, the time spent moving was divided by the sum of the time spent moving and the time spent not moving to account for instances in which the subject ant was not detected by the software. A single rotation was defined as a cumulative turn angle of 90° over a distance of 1.00cm. Turns in the opposite direction of less than 45° were ignored. The sum of both clockwise and counterclockwise rotations was used to determine rotational frequency. Trials in which the subject ant was not found for at least 95% of the recording were discarded, as were videos in which the ants appeared fatally encumbered at trial onset.

### Mechanically Evoked Biting and Mandible Opening Response (BMOR) Bioassay

To determine whether disrupting Orco-mediated olfactory signaling disrupts broadly aggression in a non-social context, individual adult minor workers were briefly anesthetized with CO_2_ before being secured with wax in a modified 200µl pipette tip such that the head and antennae were accessible. The ants were allowed to acclimate for 10 minutes before being exposed to a continuous flow of heated air alone or volatilized VU-class compounds as described above in the Volatile Orco Modulator Aggression Bioassays. A clean, ethanol washed 3.61/0.4g Von Frey hair filament (Baseline® Fold-Up™ Monofilaments Item #12-1741) was then gently brushed along the anterior portion of the ant’s head from the ventral to the dorsal side five times. Aggression was scored by six independent, blinded observers on a binary scale such that biting or attempting to bite the filament or wide opening of the mandibles (i.e. the mandibles were opened beyond parallel) either did (score of 1) or did not (score of 0) occur during the duration of the trial (Additional File 12). An aggression index was calculated by taking the average score across all observers and used for subsequent statistical analysis. Trials in which the ants had not recovered from the CO_2_ before trial onset were discarded.

### Statistical Analysis

Statistical analyses were performed using Graphpad Prism v8.0.0 (GraphPad Software, Inc). For the aggression bioassays, a two-way ANOVA was first performed followed by Holm-Sidak’s multiple comparisons test to compare NM vs. nNM aggression as well as aggression across antennal treatments. For the antennal ablation mobility controls as well as the BMOR bioassays, a Kruskal-Wallis test was performed followed by Dunn’s correction for multiple comparisons. As the volatilization mobility controls had matched samples across different time points, a repeated measures two-way ANOVA with the Geisser-Greenhouse correction for violations of sphericity was performed. For the electroantennography, linear regression analysis was used to test whether the best-fit slope differed significantly from 0 (i.e. a straight line with no dose response). The response of the hexane solvent control (i.e. 0 µg/µl of decane) was normalized to 0mV, therefore the Y-intercept was constrained to X=0, Y=0. The number of replicates for each study were as follows: Ablation Aggression Bioassays: NMs – Sham (9), U.abl (10), B.abl (6); nNMs – Sham (10), U.abl (9), B.abl (6). Mobility Controls (Ablation): Sham (29), U.abl (29), B.abl (24). Volatile Orco Modulator Aggression Bioassays: NMs – Blank (10), VUAA0 (10), VUANT1 (12), VUAA4 (10); nNMs – Blank (12), VUAA0 (11), VUANT1 (10), VUAA4 (12). Volatile Orco Modulator BMOR Bioassay: Blank (11), VUAA0 (10), VUANT1 (10), VUAA4 (10). Mobility Controls (Volatilization): Blank (8), VUAA0 (8), VUANT1 (7), VUAA4 (9). Electroantennography: Blank (5), VUAA0 (5), VUANT1 (6), VUAA4 (5). Information regarding the statistical test performed and the results from these analyses have been detailed in Additional File 3.

## List of Abbreviations

NM: Nestmate
nNM: non-nestmate
CHC: cuticular hydrocarbon
OR: odorant receptor
Orco: odorant receptor co-receptor
ORN: odorant receptor neuron (ORN)

## Declarations

### Availability of data and materials

All data generated or analyzed during this study are included in this published article (and its supplementary information files) (Additional files 3-9).

### Authors’ contributions

STF contributed to the design and conception of the work and was involved in collecting, analyzing, and interpreting the data as well as drafting and revising the manuscript. KYP and AR contributed to the conception and design of the experiments and assisted in data acquisition and analysis for the electrophysiology and aggression bioassay, respectively. IB assisted in data acquisition and analysis for the electrophysiology experiments. LJZ contributed to the design and conception of the work as well as interpreting the data and drafting and revising the manuscript. All authors read and approved the final manuscript.

## Acknowledgments

We thank the laboratories of our colleagues Dr. J. Liebig (Arizona State University) and Dr. S. L. Berger (University of Pennsylvania) for ant collections. We also thank Drs. Berger, R. Bonasio, P. Abbot, A. Carr and HW Honegger for comments on the manuscript. We lastly thank Drs. HW. Honegger, J. Slone and R. Jason Pitts and Mr. Zi Ye along with other members of the Zwiebel lab for suggestions throughout the course of this work, Dr. S. Ochieng for ant rearing and technical help and Dr. AM McAinsh for editorial assistance.

## Additional file 1 (.pptx)

Comparison of the aggression bioassay arenas (**A**) and a schematic of the volatilization bioassay schematic (**B**).

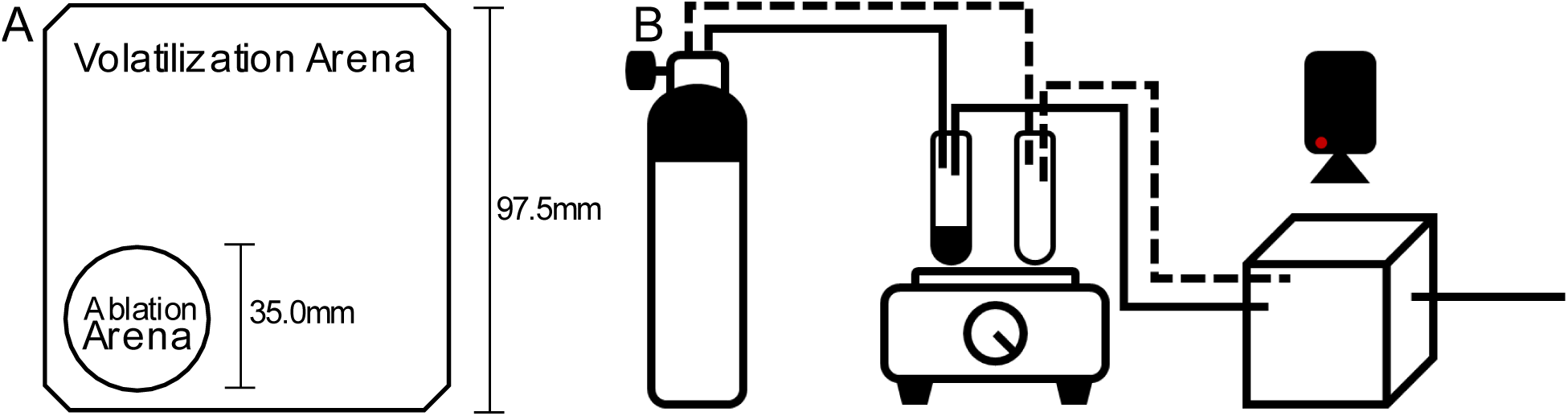

## Additional file 2 (.pptx)

Aggression (biting and wide opening of the mandibles) of individual ants in response to a mechanical stimulus from a Von Frey filament. There is no significant difference in aggression between ants exposed to either heated-air alone (Blank), VUAA0, VUANT1, or VUAA4 (Kruskal-Wallis test, N=10-11). Error bars display S.E.M.

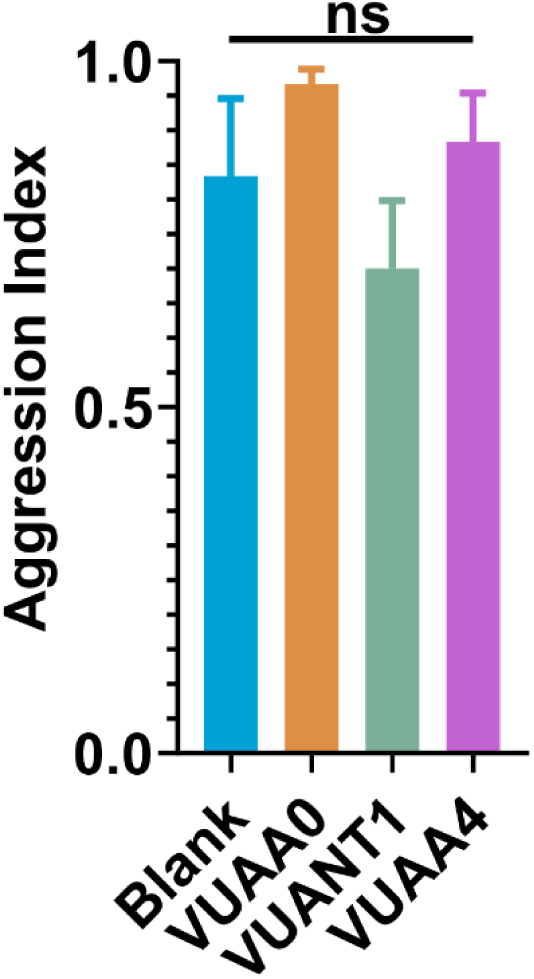

## Additional File 3 (.xls)

Summary of statistical test results.

## Additional File 4 (.xls)

Raw data for ablation aggression bioassay.

## Additional File 5 (.xls)

Raw data for mobility controls (ablation).

## Additional File 6 (.xls)

Raw data for electroantennograms.

## Additional File 7 (.xls)

Raw data for volatile Orco modulator aggression bioassay.

## Additional File 8 (.xls)

Raw data for mechanically evoked biting and mandible opening response bioassay.

## Additional File 9 (.xls)

Raw data for mobility controls (volatilization).

## Additional File 10 (.mp4)

Examples of aggression and non-aggression observed in the ablation aggression bioassay.

## Additional File 11 (.mp4)

Examples of aggression and non-aggression observed in the volatile Orco modulator aggression bioassay.

## Additional File 12 (.mp4)

Examples of aggression and non-aggression observed in the mechanically evoked biting and mandible opening response bioassay.

